# Differential dropout analysis captures biological variation in single-cell RNA sequencing data

**DOI:** 10.1101/2021.02.01.429187

**Authors:** Gerard A. Bouland, Ahmed Mahfouz, Marcel J.T. Reinders

## Abstract

Single-cell RNA sequencing data is characterized by a large number of zero counts, yet there is growing evidence that these zeros reflect biological variation rather than technical artifacts. We propose differential dropout analysis (DDA) to identify the effects of biological variation in single-cell RNA sequencing data. Using 16 publicly available and simulated datasets, we show that DDA accurately detects biological variation and can assess the relative abundance of transcripts more robustly than methods relying on counts. DDA is available at https://github.com/gbouland/DDA.

## Main

Single cell RNA sequencing (scRNAseq) data is highly sparse, and the common belief is that the zero values are primarily caused by technical artifacts (i.e. dropouts). Although more zeros are observed in scRNAseq data than expected, these can largely be explained by biological rather than technical factors^1^. Also, the amount of zeros in scRNAseq is in line with distributional models of molecule sampling counts^2^. Methods that utilize dropout patterns for feature selection^3,4^ and cell type clustering^5^ have recently been developed and perform better or comparable with methods relying on highly variable genes. For instance, Qiu^5^ binarized scRNAseq count data, where dropouts remain zero and every non-zero value was assigned a one. With this binary representation, cell type clusters were identified based on co-occurrence of transcripts. Yet, it is not clear whether dropouts also reflect differences across distinct biological cell populations. Therefore, we investigated whether biological differences across cell population can be identified using differential dropout analysis (DDA), rather than the commonly used differential expression analysis (DEA). Instead of relying on changes in the expression value of genes across cell populations, which can be sparse and are subject to pre-processing steps, we analyzed the binary dropout patterns across biological distinct cell populations, i.e. are there more (or less) dropouts for a gene in condition *A* compared to condition *B*.

As proof of concept, we performed DDA with a simple logistic regression on binarized expression profiles from 16 scRNAseq datasets (662,825 cells in total, Supplementary Table 1). We compared the results of DDA with those of differently expressed genes (DEGs) detected using the commonly used Wilcoxon Rank Sum test, which is also top ranked for single cell analyses^6,7^. We tested each gene using both DDA and DEA for differences between conditions (6 datasets), cell-types (6 datasets) and normal-versus cancerous tissues (4 datasets, Fig. 1a). Across all datasets, a total of 96,275 significant genes (P_FDR_ ≤ 0.05) were identified with either DDA (92,381 genes) or DEA (91,521 genes). Of these, 87,627 were identified by both tests, resulting in a Jaccard index of 0.91. This high degree of agreement is also reflected in each individual dataset (median = 0.92, minimum = 0.76, and maximum = 0.99). We did not use a log fold-change (logFC) or log odds-ratio (logOR) threshold, as for each dataset and comparison different thresholds are appropriate. In all datasets, the logFC and logOR were significantly (spearman) correlated (median(ρ) = 0.90, minimum(ρ) = 0.49, and maximum(ρ) = 0.98, P ≤ 5 × 10^−100^). The three datasets with the lowest correlation coefficient between logOR and logFC (ρ ≤ 0.62) were datasets generated using the Smart-seq protocol (Supplementary Table 1). Across the datasets, we observed an average increase of 1.80 in logOR (median = 1.70, Q_1_ = 1.59, Q_3_ = 2.10), for every increase in logFC (see Fig. 1b for the *cancer atlas (2)* dataset^8^). The high degree of agreement of detected genes shows that DDA performs on par with the Wilcoxon Rank Sum test and the strong correlation of the logFC and logOR across all datasets shows that the results can be interpreted in a similar way.

**Figure 1.**
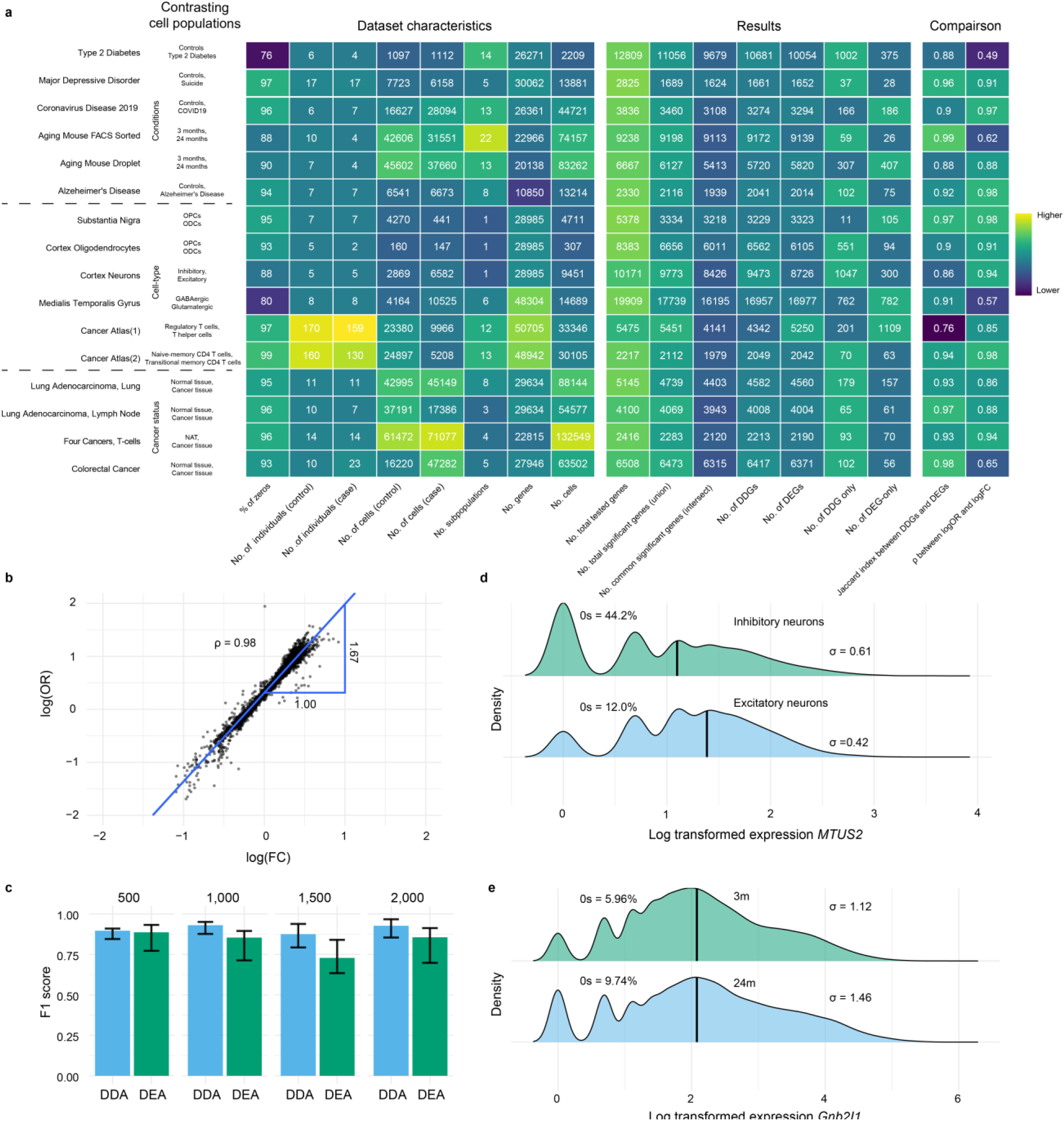
**a**, Heatmap of the dataset characteristics, general overview of the results of both DDA and DEA per dataset, and a comparison of the results. The rows represent the datasets, the first column shows the cell populations that were used as contrast for testing. **b**,Plot of the logOR and logFC of the Cancer Atlas (2) dataset. The x-axis represent the logFCs of each tested gene, and the y-axis represent the logORs for the same genes. The blue lines shows the linear association between the logFC and logOR. The Spearman’s rank correlation coefficient (*ρ*) is also shown in the plot. **c** Barplots of the F-score of DDA based on simulated data. Numbers above barplots shows the number of cells that were generated for the repsective simulations simulation. Height of bar is defined the median value from 25 simulations, error bars are defined by the first and thrid quartile. **d** Two density plots of *MTUS2* from cortex neuron dataset. The top plot shows the density of *MTUS2* in inhibitory neurons and the bottom plot shows the density of *MTUS2* in excitatory neurons. **e** Two density plots of *Gnb2I1* from the aging mouse atlas droplet dataset. The top plot shows the density of *Gnb2I1* in 3-month-old mice and the bottom plot shows the density of *Gnb2I1 in* 24-month-old mice. Both **d** and **e** are supported with fraction of zeros and variance of each cell population.

To compare the performance of DDA and DEA in a controlled manner, we simulated scRNAseq data with muscat^9^ using the provided dataset as reference^10^. We generated scRNAseq data with varying number of cells and 25% of differentially expressed genes. With these simulated data, DEA detected consistently more genes, both the true positive rate (TPR) and false positive rate (FPR) were higher than the TPR and FPR of DDA (Supplementary Fig. 1a, Supplementary Fig. 1b). DDA, however, had a higher precision (PPV, Supplementary Fig. 1c) and was more accurate (F1 score, Fig. 1c).

Despite the observed association between mean expression and dropout rate, which has been previously described^2^, and similar performance of the two tests, there were 4,754 and 3,894 genes uniquely identified using DDA and DEA, respectively, across all datasets. To better understand these differences, we highlighted two differential dropout genes (DDGs). In the *cortex dataset*^11^, *MTUS2* had significantly less dropouts in excitatory neurons (logOR = 1.30, P_FDR|DDA_ = 5.27 × 10^−115^, Fig. 1d) compared to inhibitory neurons, while the median expression levels were not significantly different (logFC = −1.70 × 10^−3^, P_FDR|DEA_ = 5.70 × 10^−2^), implying additional high ranked expressions for every additional zero. In the *aging mouse atlas droplet* dataset^12^, *Gnb2l1* had significantly less dropouts in the 3-month-old mice (logOR = −0.67, P_FDR|DDA_ = 1.81 × 10^−122^, Fig. 1e) compared to the 24-month-old mice, while again the median expression levels were not significantly different (logFC = 3.15 × 10^−3^, P_FDR|DEA_ = 6.97 × 10^−1^). These examples show that differences in variance between contrasting cell populations can interfere with the association between dropout rate and mean expression, resulting in disparities between DDA and DEA.

To exclude that the differentially behaving genes between DDA and DEA associate with a specific technological or biological process, we investigated whether there were genes repeatedly detected by a one of the two methods. In most cases, genes that were identified as DDG-only (or DEG-only) were found within a single dataset (Supplementary Fig. 2a, Supplementary Fig. 2b), suggesting the absence of a driving process for them.

To validate the DDA results, we compared the results of the *Alzheimer’s Disease* (AD) dataset^13^ (entorhinal cortex) with DEA analysis performed on a bulk RNAseq AD dataset^14^, comprised of samples from the fusiform gyrus. For genes measured in both, the scRNAseq dataset and the bulk dataset (N = 2,228), the majority of DDG-only (60.7%) were also differentially expressed in bulk (Fig. 2a). The logOR of the single cell analysis was also significantly correlated with the logFC of the bulk analysis (ρ = 0.39, P = 9.70 × 10^−79^, Fig. 2b). Similarly, in a second dataset^15^, 65.5% of the DDG-only genes were differentially expressed in bulk samples from the frontal cortex, temporal cortex and hippocampal formation (Supplementary Fig 3). The high correlation between the logOR and the logFC of a bulk dataset, despite being from different brain region, and that the majority of DDG-only were still detected in bulk, further confirms that DDA can be used to detect differentially abundant genes.

**Figure 2.**
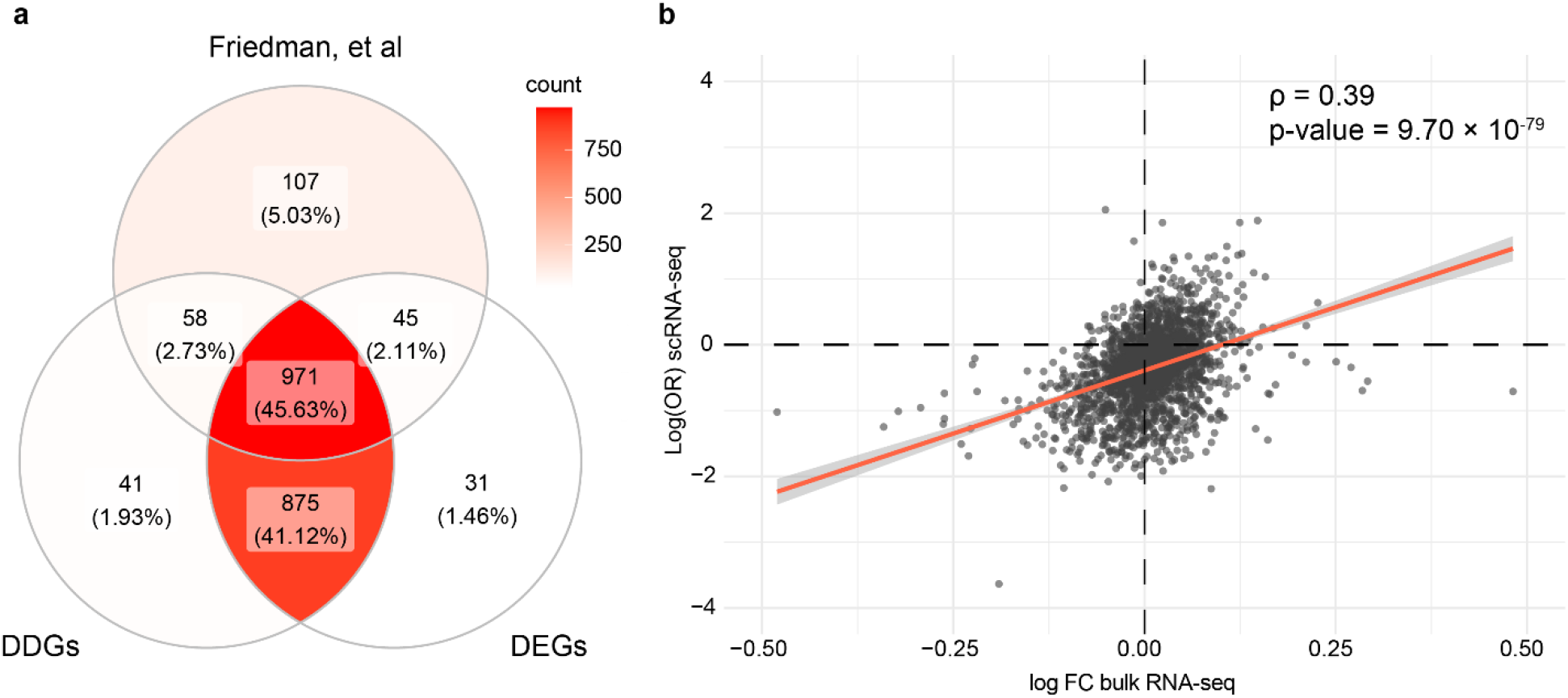
**a**,Venn diagram of genes detected (P_FDR_ ≤ 0.05) in a bulk AD dataset (Friedman, et al), in the single cell AD dataset with DDA (DDGs) and with DEA (DDGs). Each section shows the number and percentage of genes belonging to that section. **b** Plot of the logFC from the AD bulk dataset (x-axis) and the logOR from the single cell AD dataset (y-axis). The red line represents the linear association between the bulk logFC and logOR. The Spearman’s rank correlation coefficient (ρ) and corresponding association p-value are also shown. Outlier genes (n = 9, **b**) were removed from the plots.

To test the binarization scheme, we performed a DDA on the AD dataset for binary profiles generated with different thresholds for binarization (thresholds ranging from one to ten counts). Naturally, for every increase in the threshold, the number of genes with zero measurements across all cells increased, resulting in a decreasing number of tested and significant genes (Supplementary Fig 4a, Supplementary Fig 4b). With higher thresholds, we found a decrease in correlation of the logORs from DDA with the logFC from DEA (Supplementary Fig 4c). These results show that the default binarization scheme where dropouts remain zero and every non-zero value is assigned a one, is indeed appropriate.

Finally, a runtime benchmark comparing DDA with all DE methods available in Seurat revealed that DDA is among the most time efficient methods for detecting differentially abundant genes (Supplementary Fig 5).

Altogether, our results show that dropout patterns across cell populations represent biological variation and can be used as measure of relative abundance of transcripts. Across 16 datasets and a variety of contrasting cell populations (disease vs healthy, cell-types, and cancer status), DDA detected biologically relevant genes that were missed by DEA. While the performance of DDA and DEA on real data is largely comparable, with a known ground truth, DDA performed better than DEA on simulated data. Additionally, DDA benefits reproducibility and is more robust than DEA, since the only pre-processing step required for DDA is the binarization of counts. In contrast, DEA requires normalization and transformation of counts, where an analyst can choose from an excess of equally valid methods^16^. Performing DDA on datasets generated using the Smart-seq protocol should be approached with more caution: although the agreement of detected genes between DDA and DEA was high, we observed the lowest correlation between the logOR and logFC for these datasets.

DDA is a normalization-free, time efficient and accurate alternative to DEA for which we see three potential use cases. First, DDA could be performed in isolation as a fast and accurate alternative to DEA. Besides logistic regression, alternative tests such as the Fisher’s exact test or the chi-squared test could also be explored. However, logistic regression is more versatile as it allows to adjust for covariates. Second, DDA could be performed in addition to DEA to identify more genes. Finally, DDA could be used to validate pre-processing, normalization and DEA as a big discrepancy between the DDGs and DEGs could indicate an aberration in the DEA results.

## Methods

### Single cell RNA datasets

In total, 16 scRNAseq dataset (14 human and 2 mouse) were used to investigate the utility and biological relevance of binarized expression profiles of genes (Supplementary Table 1). All datasets had pre-annotated cell types and conditions. From the corresponding references, un-normalized count matrices were acquired, and only annotated cells were kept for further analysis. For each dataset, we extracted the annotated cell-type, patient ID, and to which of contrasting cell population the cell belonged from the included meta data. This was slightly different for the aging mouse atlases^12^ and cancer atlas^8^. For the aging mouse atlases, instead of annotated cell-types we retrieved the tissue names. For the cancer atlas, the contrasting cell populations were defined by cell-type, so we retrieved the tissue and the cancer-type for each cell. Each dataset was separately pre-processed. For the differential dropout analysis (DDA), the count matrices were transformed to a binary representation, where dropouts remain zero and every non-zero value was assigned a one. For the differential expression analysis (DEA), each count matrix was log-normalized using Seurat 3.2.2^6^, such that: 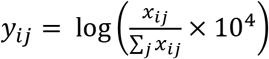, where *x*_*ij*_ and *y*_*ij*_ are the raw and normalized values for every gene *i* in every cell *j*, respectively. This normalizes the feature expression measurements for each cell by the total expression, multiplies this by a scale factor (10,000 by default), and log-transforms the result. The cancer atlas was already normalized, as it was a merger of multiple datasets.

### Statistical analysis

Association p-values were corrected for multiple tests with the Benjamini-Hochberg procedure and significance was assumed at an adjusted P-value of P_FDR_ ≤ 0.05. Spearman’s rank correlation coefficient and the associated p-values were calculated using the `cor.test` function in R v4.0.2.

### Differential expression analysis

DEA was performed using the Wilcoxon Rank Sum test using the FindMarkers function in Seurat 3.2.2^6^. Note that the Wilcoxon Rank Sum test from Seurat takes into account zero measurements, and handles them as ties between contrasting cell populations. Genes coding for ribosomal proteins were excluded and we only tested genes that were expressed in at least 10% of the cells in either of the respective groups of interest. This is the default option in the FindMarkers function and speeds up testing by ignoring infrequently expressed genes. Genes coding for ribosomal proteins were excluded. Association p-values were corrected for multiple tests.

### Differential dropout analysis

DDA was performed using logistic regression with the `glm()` function in R v4.0.2. All genes that were tested with DEA were also tested with DDA. The resulting association p-values were corrected for multiple tests.

### DDA – DEA comparison

For the comparison between DDA and DEA, we investigated agreement and disagreement between detected genes and the linear association between the logOR and logFC. Agreement was calculated by the Jaccard index, i.e. number of genes that both tests commonly detected, divided by the total number of genes that were detected. Agreement was calculated on the combination of all datasets and for each individual dataset. The disagreement was investigated by means of inspecting characteristics of DDGs-only and DEGs-only. DDGs-only were defined as genes that were detected (P_FDR_ ≤ 0.05) by DDA and were not detected (P_FDR_ > 0.05) by DEA. Conversely, DEGs-only were defined as genes that were detected (P_FDR_≤ 0.05) by DEA and were not detected (P_FDR_ > 0.05) by DDA. The Spearman’s rank correlation coefficients between the logOR and logFC were calculated with the estimates of all tested genes of the respective datasets. The scale differences for every dataset, between logOR and logFC, were calculated with a linear model on the estimates of all tested genes of the respective datasets, using the `lm` function in R v4.0.2. The logOR was specified as outcome variable and the logFC as predictor variable. The resulting slopes were interpreted as scale differences between the logOR and logFC

### Simulation

Data was simulated with muscat 1.2.1^9^. The provided dataset^10^ was used as reference. 100 simulated datasets were generated with varying sample sizes (500 cells, 1,000 cells, 1,500 cells and 2,000 cells), 25 simulations per sample size. For each simulation 1,000 genes were generated of which 25% were differently expressed between two groups of equal size. For both tests we calculated the True Positive Rate (TPR), False Positive Rate (FPR), Positive Predictive Value (PVV) and accuracy (F1-score) per simulation.

### Running times

The runtime benchmark was performed with simulated data from muscat^9^. Runtimes of DDA and all DE methods (wilcox, bimod, roc, t, negbinom, poisson, LR, MAST and DESeq2) included in Seurat were measured based on 500 genes and 1,000 cells up to 10,000 cells with increments of 1,000. Runtime was measured with the `system.time` function in R.

### Validation with existing bulk RNA-seq data

The AD bulk RNA-seq datasets were acquired from Gemma^17^. The first dataset from Friedman, et al^14^ consisted of 33 controls (CT) and 84 samples from individuals diagnosed with Alzheimer’s Disease (AD) collected from the fusiform gyrus. This dataset was reprocessed by Gemma and no batch effects were present. For the differential expression analysis in bulk we used the Wilcoxon Rank Sum test from limma v3.44.3^18^. In total, 2,228 genes were tested for differential expression, as these genes were also included in the scRNAseq AD dataset^13^ analysis. The second dataset from Hokama, et al^15^ consisted of 47 controls and 32 AD samples. The samples originated from the frontal cortex (N_CT_ = 18, N_AD_ = 15), temporal cortex (N_CT_ = 19, N_AD_ = 10) and hippocampal formation (N_CT_ = 10, N_AD_ = 7). The data was reprocessed and batch corrected by Gemma. For the differential expression analysis no distinction was made between brain regions. In total, 2,001 genes were tested for differential expression. All resulting association p-values were corrected for multiple tests. For validation, the significant DDGs and DEGs from the scRNAseq AD dataset analyses were compared with the significantly differentially expressed genes from the bulk analyses. Venn diagrams were plotted with ggVennDiagram v0.3^19^. Correlations were calculated between the logOR and logFC of the single-cell analysis with the bulk logFC.

## Availability of data and materials

The datasets used and prepared for this study can be downloaded from Zenodo (http://doi.org/10.5281/zenodo.4487320). The results are also made available in an interactive shiny dashboard: http://insyprojects.ewi.tudelft.nl:5000/DifferentialDropoutAnalysis/. The scripts, functions and source data for the figures are available at the Github repository: https://github.com/gbouland/differential-dropout-analysis, including two vignettes describing a DDA starting from a Seurat object and a raw count matrix. DDA is also implemented in an R-package and is available at: https://github.com/gbouland/DDA.

## Acknowledgements

This research was supported by an NWO Gravitation project: BRAINSCAPES: A Roadmap from Neurogenetics to Neurobiology (NWO: 024.004.012)

## Author contributions

GAB, AM, and MJTR conceived the study and designed the experiments. GAB performed all experiments and drafted the manuscript. GAB, AM, and MJTR reviewed and approved the manuscript.

