## Supplementary Figures for "Differential dropout analysis captures biological variation in single-cell RNA sequencing data"

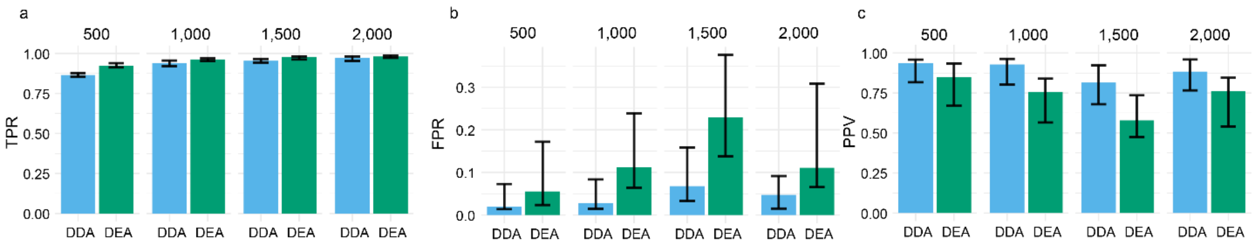


***Supplementary Figure 1*** Barplots of evaluation of DDA and DEA on simulated data **a,** True positive rate (TPR). **b,** false positive rate (FPR). **c,** and positive predictive value (PPV). Numbers above barplots shows the number of cells that were generated for the respective simulations simulation. Height of bar is defined the median value from 25 simulations, error bars are defined by the first and third quartile.

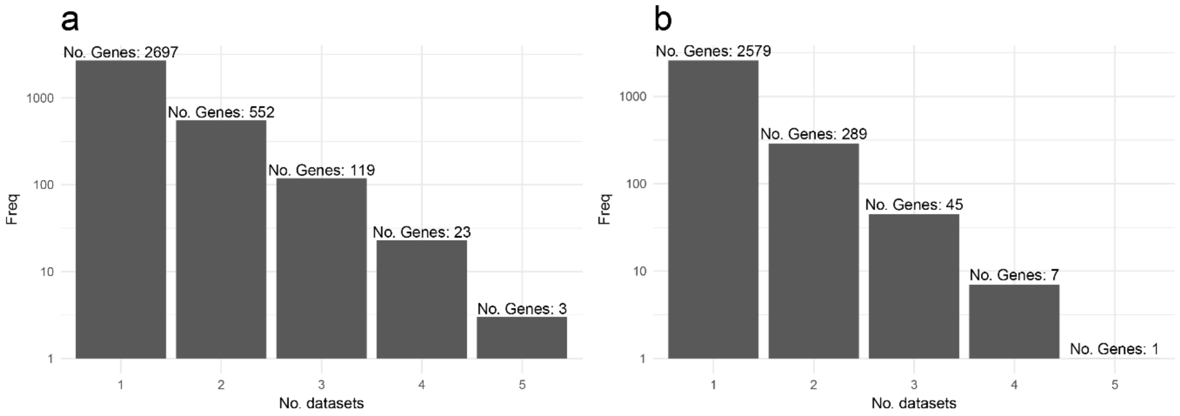


***Supplementary figure 2 a,*** Bar plot of the number of genes that were identified as DDG-only in *i* datasets. The x-axis represent the number of datasets and the y-axis represent the number of genes that were found with that many datasets. For instance, 2740 genes were detected with a single dataset and 3 genes were detected with 5 datasets. ***b,*** Bar plot of the number of genes that were repeatedly identified as DEG-only. The x-axis represent the number of datasets and the y-axis represent the number of genes that were found with that many datasets.

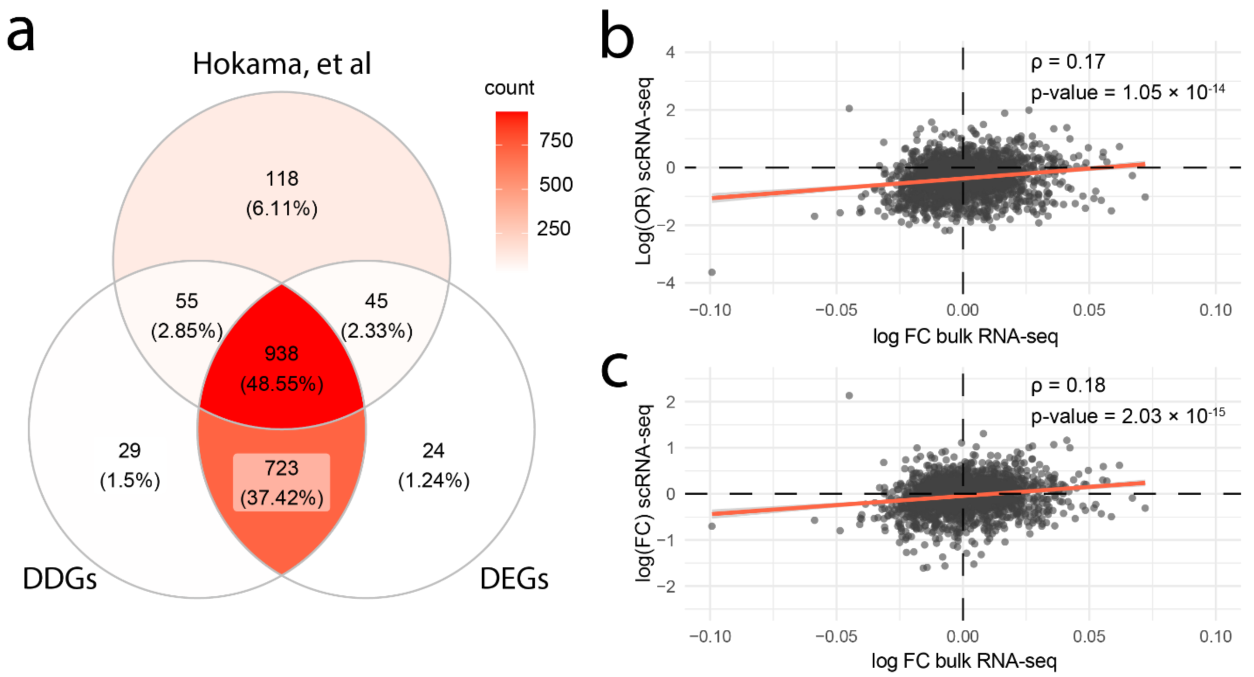


***Supplementary figure 3 a,*** Venn diagram of genes detected ($P_{FDR} \leq 0.05$) in bulk AD dataset (Hokama, et al), in the single cell AD dataset with DDA and with DEA. Each section shows the number and percentage of genes belonging to that section. **b,** Plot of the logFC from the AD bulk dataset (x-axis) and the logOR from the single cell AD dataset (y-axis). The red line represents the linear association between the bulk logFC and single cell logOR**.** The Spearman's rank correlation coefficient (rho) and corresponding association p-value are also shown. **c,** Plot of the logFC from the AD bulk dataset (x-axis) and the logFC from the single cell AD dataset (y-axis). The red line represents the linear association between the bulk logFC and single cell logFC**.** The Spearman's rank correlation coefficient and corresponding association p-value are also shown.

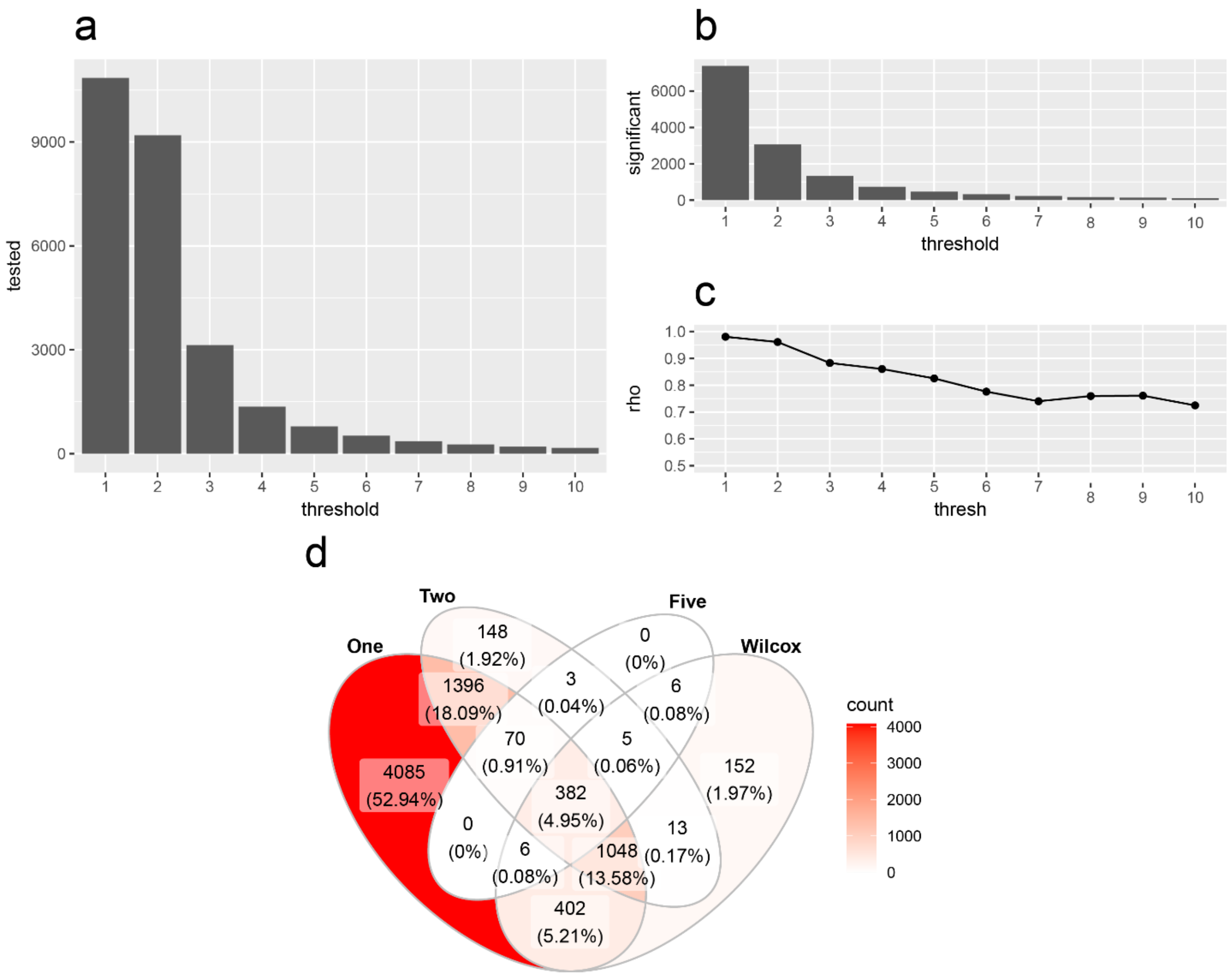


***Supplementary figure 4 a,*** Bar plot of total number of tested genes at each binarization threshold. The x-axis represents the binarization threshold and the y-axis represent the number of genes that were tested with that threshold. ***b,*** Bar plot of total number of significant genes ($P_{FDR} \leq 0.05$) at each binarization threshold. The x-axis represents the binarization threshold and the y-axis represent the number of genes that were significant with that threshold. ***c,*** Pearson's correlation coefficient of the logOR of the DDA with different thresholds with the logFC of the DEA. The x-axis represents the binarization threshold and the y-axis represents The Spearman's rank correlation coefficient with that threshold. ***d,*** Venn diagram of detected genes ($P_{FDR} \leq 0.05$) with a threshold of 1, 2, 5 and DEA (wilcox). Each section shows the number and percentage of genes belonging to that section.

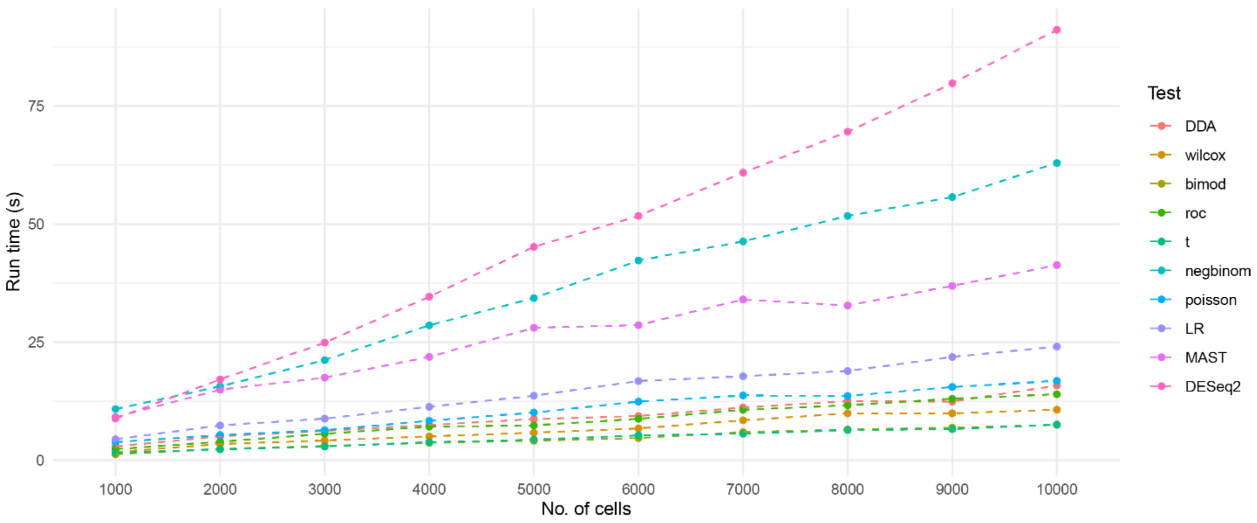


***Supplementary Figure 5*** Results of benchmark in which run time was evaluated. The x-axis represents the number of cells for the respective run, the y-axis represents the run time in seconds. All runs were performed with 500 genes.
